# Endometrial Gap Junction Expression - Early Indicators of Endometriosis and Integral to Invasiveness

**DOI:** 10.1101/2021.01.25.428135

**Authors:** Chen-Wei Chen, Jeffery Chavez, Li-Ling Lin, Chiou-Miin Wang, Ya-Ting Hsu, Matthew J. Hart, Jianhua Ruan, Laurel Gillette, Richard O. Burney, Robert S. Schenken, Randal D. Robinson, Maria Gaczynska, Pawel Osmulski, Nameer B. Kirma, Bruce J. Nicholson

**Author notes:** Equal contributions as first authors (Chen-Wei Chen, Jeffery Chavez). Co-corresponding authors, (NBK) (BJN).

## Abstract

Endometriosis is an invasive disease, and a leading cause of pain, infertility and disability among women, with an incidence 10 fold that of cancer. A more complete understanding of disease pathogenesis is essential for the development of non-surgical diagnostic assays and non-hormonal therapeutics. Avoidance of immune clearance and implantation of endometrial tissue on peritoneal surfaces are features of endometriosis lesion formation that overlap with cancer metastasis. Connexins, and the gap junctions they form, have been implicated in cancer progression, and may be associated endometriosis pathophysiology. Single cell transcriptomic profiling of endometrial epithelial and stromal cells from women with endometriosis reveals a striking and progressive shift in expression of connexins and related regulatory and junctional genes. We demonstrate that gap junction coupling between endometrial cells and the peritoneal mesothelium is dramatically induced, specifically in endometriosis patients, and is required for invasion by inducing breakdown of the mesothelial barrier function.

## INTRODUCTION

Endometriosis is a chronic inflammatory disease affecting 6-10% of reproductive age women (Eskenazi and Warner,1997). Characterized by the presence of endometrial tissue in extrauterine locations including the pelvic peritoneum, ovary and bowel surface, endometriosis is diagnosed in 35-50% of women with pelvic pain and up to 50% of women with unexplained infertility (Rogers et.al., 2009). At an estimated annual cost of $12,000 per patient in terms of diagnosis and treatment, and adding in the significant loss of productivity, endometriosis care entails significant socioeconomic burden for both individual patients and healthcare systems estimated to cost $80 billion per year for the US alone (Soliman et.al., 2016). In the absence of a biomarker, laparoscopic surgery remains the gold standard for diagnosis. The requirement for invasive surgery, which fails to confirm endometriosis almost half the time (Mettler et.al., 2003) contributes to an average latency of 6.7 years from onset of symptoms to definitive diagnosis (Bontempo and Mikesell, 2020), and results in 68% of women suffering from endometriosis being incorrectly diagnosed (Hudelist et.al., 2012). Diagnostic delay allows time for disease progression, and potentially worsens sequelae and prognosis. An improved understanding disease etiology is critical to developing new diagnostics and therapies.

The original, and still most widely accepted, model for the pathogenesis of endometriosis is retrograde menstruation, in which sloughed endometrial tissue during menses traverses the fallopian tubes and forms invasive lesions within the peritoneal cavity (Sampson, 1927). Peritoneal origins have also been proposed (Mismer et.al., 2004; Sasson and Taylor, 2008), but the preponderance of evidence still favors endometrial origins (reviewed in Burney and Giudice, 2012). Abundant evidence supports molecular differences in the eutopic endometrium of women with and without endometriosis [Burney et.al., 2007; Rogers et. al., 2009; Ulukus et. al., 2006; Yu et.al.,2014] suggesting enhanced survival (Jones et.al., 1998) and invasive potential (Lucidi et.al., 2005) of endometrium from affected women. These innate or acquired molecular features distinguish women with endometriosis, and may predispose endometrial tissue to invade and form lesions when transported into the pelvic cavity and distinguish women destined to develop endometriosis [Hastings and Fazleabas, 2006, Tamaresis et.al., 2014]. Since retrograde menstruation is estimated to occur in 76-90% of women, but only 6-10% develop endometriosis (Burney and Giudice, 2012) the interaction of displaced endometrium and the peritoneum is a key differentiating step in disease pathogenesis. Peritoneal factors also influence disease progression, including the hormonal environment (Parente Barbosa et.al., 2011), oxidative stress and inflammation (Augoulea et.al., 2012) or decreased immune clearance (Oosterlynck et.al., 1991), although this could be due to changes in the endometrial cells themselves (Somigliana et.al., 1996). This has fueled the debate over whether endometriosis originates from changes in the uterus that predispose the cells to lesion formation (the “seed” model) or if it is more a property of a receptive peritoneal environment (the “soil” hypothesis).

To understand the molecular underpinnings of endometrial-peritoneal interaction in lesion formation, we have focused on a class of proteins that has been implicated in tissue invasion, infertility and inflammation in other contexts, but incompletely explored in the pathophysiology of endometriosis. Gap junctions, composed of connexin (Cx) proteins encoded by a family of 21 GJ (A-D) genes, mediate direct contact and communication between most cells of the body via exchange of ions as well as metabolites and signaling molecules <1kD (Goldberg et.al, 1999; Weber et.al, 2004). Gap junctions have been shown to be <u>essential to many invasive processes</u>, both normal (e.g. blastocyst implantation – Grummer et.al., 1996) and pathogenic (e.g. metastasis). Increased Cx43 and Cx26 expression and gap junction intercellular communication (GJIC) have been associated with metastasis and poor prognosis in breast (Kanczuga-Koda et.al., 2006; Naoi et.al. 2007; Stoletov et.al., 2013) and prostate cancers (Zhang et.al., 2014; Lamiche et.al., 2012), and elevated Cx26 with melanoma metastases (Ito et.al., 2000). Connexins exert their effects both during intravasation and extravasation (as cells enter and leave the circulation) [el Sabban and Pauli, 1991, 1994; Ito et.al, 2000, Naoi et.al, 2007], as well as establishment of distant lesions by forming heterotypic GJIC (Lamiche et.al., 2012; Stoletov, et.al., 2013; Hong et.al, 2015; Chen et.al., 2016). In a global screen of cervical squamous carcinoma, Cx43 emerged as one of three genes (along with PDGFRA2 and CAV-1) central to cancer invasion and metastasis (Cheng et.al., 2015).

<u>Gap junctions are also critical</u> for a number of steps in human fertility (<u>reviewed in Winterhager and Kidder, 2015</u>). Endometrial gap junctions, comprised of Cx43, are essential for decidualization [Kaushik et.al., 2020], blastocyst implantation [Grummer et.al.,1996: Diao et.al., 2013] and vascularization and endometrial development during pregnancy [Laws et.al., 2008]. <u>Connexins have also been linked to induction of inflammatory processes</u> that either inhibit tissue repair (Willebrords et.al., 2016), such as wound healing of the skin (Montgomery et.al., 2018) and cornea (Ormonde et.al., 2012) processes thought to be driven by ATP release through connexin hemichannels (Mugisho et.al., 2018).

With regard to endometriosis, prior immunohistochemistry studies demonstrated a shift in Cx expression of endometrial epithelial cells (EECs) from primarily Cx26, with some Cx32 (*GJB1*), in the uterus to Cx43 in peritoneal (ectopic) endometriotic lesions (Regidor et.al., 1997). By contrast, endometrial stromal cells (ESCs) show predominantly Cx43 expression in both eutopic and ectopic locations, although at reduced levels in endometriosis patients (Yu et.al., 2014). This modified expression profile of connexins seen in ectopic lesions of women with endometriosis was recapitulated in eutopic endometrial tissue in an endometriosis model in baboons (Winterhager et.al., 2009), suggesting that early changes in the endometrium might predispose refluxed menstrual endometrial tissue within the pelvic cavity for invasiveness (Guo et.al., 2004; Lucidi et.al., 2005;).

Our objective in this study was to characterize the contribution of Cxs within the endometrial compartment to its invasive potential in the peritoneum, an essential process for endometriosis lesion formation. We have examined the connexin gene expression profiles in both endometrial stromal (ESC) and epithelial (EEC) cell compartments, using single cell analysis of tissue obtained by endometrial pipelle biopsy from women with and without endometriosis. We also performed functional studies by assessing GJIC in both homotypic ESC and EEC cultures, as well as heterotypic co-cultures with peritoneal mesothelial cells (PMCs). We finally assessed the role of the primary gap junction (*GJ*) gene product, Cx43, in the trans-mesothelial invasive process by ESCs and EECs that is essential for endometriosis lesion formation. These studies provide unique insights into the endometrial-peritoneal interaction in lesion formation, and represent the first single cell transcriptomic analysis of connexins and gap junctions in endometriosis.

## RESULTS

### Expression profiles of Intercellular Interaction Genes in Endometrial Stromal and Epithelial Cells from Women with and without Endometriosis

While studies have addressed different aspects of endometriosis, from changes in hormonal responses and other contributors to infertility to inflammatory responses caused by peritoneal lesions, and some even on lesion formation, none have focused on what common themes may tie all these symptoms together. Given the links, described above, of connexins and gap junctions to each of these processes, we examined their expression patterns, and those of related genes involved in intercellular interactions and their regulation in the endometrial lining of normal and endometriosis patients by analysis of freshly isolated primary cells (i.e. P0). Specifically, the gene panel included: <u>13 of the total of 21 Cx genes</u>, including the major expressers in the endometrium, Cx43 (*GJA1*), 26 (*GJB2*) and 32 (*GJB1*), and at least one from each Cx class (*A*, *B* and *C*); <u>adhesion proteins</u> [N-cadherin (*CDH2*) and Ep-CAM], <u>cytoplasmic proteins associated with junctional complexes</u> [Drebin (*DBN1*), β-catenin (*CTNNB1*), caveolin (*CAV1), TJAP and ZO1 and 2 (TJP1 and 2*)]; <u>regulatory components of junctions</u>, such as <u>transcription factors</u> [Snail 1 (*SNAI1*)] and <u>kinases</u> (e.g. *PKA, PKC*, and *MAPK* isoforms, *CK1*, *SRC, CDK1*); components of the <u>extracellular matrix</u> (*MME, NOV*); housekeeping (*UBB, GAPDH, ACTIN*, and *GUSB*) and cell marker (Vimentin (VIM) and cytokeratin (KRT18)) genes. The genes, in the array, in order of presentation in the heat map, are provided in **Table S1**. Those shaded in grey were control genes fpr data normalization and are not presented in the arrays.

To optimally understand how changes in patterns of expression might contribute to disease etiology, we separated the two major endometrial cell types [epithelial or glandular (EECs) and stromal (ESCs)], the purity of which was demonstrated to be ~95% by immunocytochemistry (**Fig. S1**). To further maximize detection of micro-heterogeneities in expression patterns, we used the Fluidigm C1 single-cell capture system followed by the Biomark microfluidic PCR to examine expression at the single cell level within each population. Two normal subjects and 6 endometriosis patients (one in early stage (I-II), and the others later stage (III-IV) endometriosis) were examined, with samples taken from proliferative, early and late secretory menstrual phases, plus one non-cycling patient on birth control (**Table 1**). Normalized expression results for each cell type from each patient are displayed as a heat map, with each row representing a gene, and each column a cell (**Fig. 1A, B**). Distinct patterns of expression in stromal (ESCs) and epithelial cells (EECs) were evident, which in this patient sampling clearly distinguished normal from endometriosis samples, and even showed a gradual transition as disease progressed from early to late stages (**Fig. 1A, B**).

**TABLE 1:**
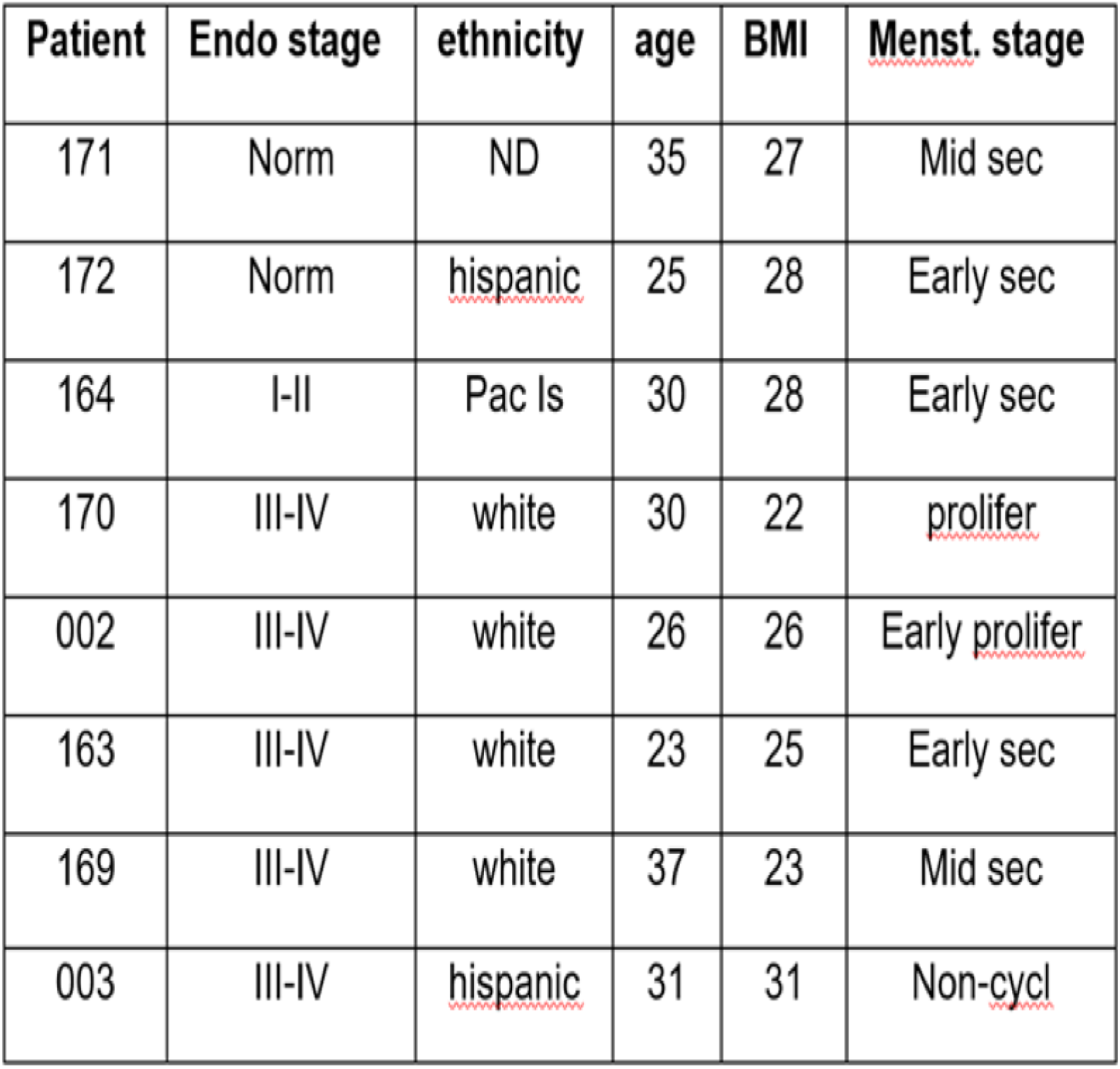
Patient database

**Figure 1:**
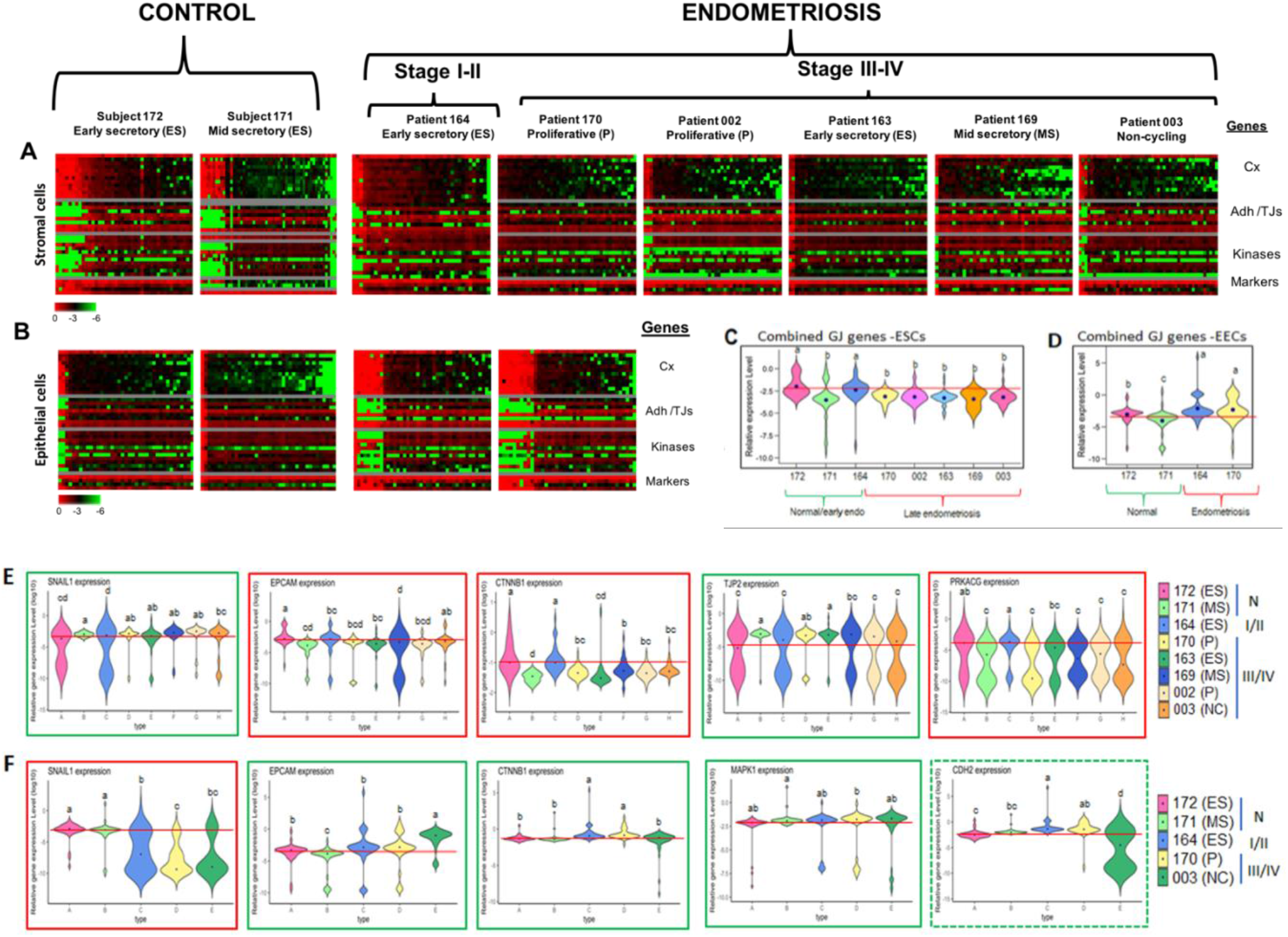
Single cell gene expression profiles show a consistent, but opposite, shift in Gap Junction (*GJ*) gene expression in endometrial stromal (ESCs) and epithelial cells (EECs) with endometriosis progression. (**A-B**) Heat maps of single cell (columns) expression of a panel of genes, plotted in descending order of Cx gene expression (rows) involved in intercellular interactions and their regulation in ESCs (**A**) and EECs (**B**) isolated from 2 control and 6 endometriosis patients, with samples taken at different disease stages and at different phases of the menstrual cycle, as indicated. Due to poor yields or their adhesive nature preventing single cell isolation for analysis, EECs could not be analyzed for all endometriosis patients. Most notable were changes in the abundance of cells with high levels of expression of all *GJ* genes (top third of each map), which steadily decreased with disease progression in ESCs and increased in EECs. A full list of genes in order (top to bottom) in **Table S1. (C-F**) Violin plots showing the distribution of RNA expression levels (log scale) for selected genes, or groups of genes, in stromal (C, E) and epithelial cells (D, F) derived from each of the patients studied (indicated by numbers on the X-axis). C and D show the aggregate of combined *GJ* gene expression, while E and F show 6 genes with the most consistent changes with endometriosis. Green outlines indicate increased expression and red outlines decreased expression in endometriosis patients compared to control (EECs) or control and early endometriosis, excluding the sample taken during mid-secretory phase of the menstrual cycle (ESCs), indicated by the red horizontal line in each plot. Horizontal bulges in each plot indicate larger numbers of cells with that expression level. The black dot in the middle of the plot indicates mean expression. Significance was determined by Duncan’s multiple range test and statistical differences (P<0.05) between samples, as indicated by letters (e.g. a vs. b; b vs. d). In addition, ‘a’ indicates the highest mean expression and ‘d’ indicates the lowest mean expression value for each factor (i.e. a>b>c>d). Violin plots of the remaining genes are shown in **Fig. S2**.

#### (a) Gap Junction genes

With respect to genes encoding connexins (*GJ* genes – located at the top of the maps), a subset of ESCs (~25% of total analyzed cell population) showed high relative expression of all *GJ* genes in normal subjects, which diminished somewhat in early stage (I/II) endometriosis samples, and drops to less than 10% in late stage (III/IV) endometriosis, regardless of the menstrual phase from which the sample is collected (**Fig. 1A**). Even though single EEC cell preparations could not be made from all patients, it was clear that the pattern was completely reversed compared to ESCs, with low expression of *GJ* genes in normal subjects (5% of cells show higher expression), which progressively increases in early (I/II) and late stage (III/IV) endometriosis where ~25% of cells show high expression (**Fig. 1B**). It is striking that this is true for all *GJ* genes, and not just the most abundantly expressed, like *GJA1*(Cx43). The combined expression levels of all *GJ* genes is illustrated in Violin plots for ESCs (**Fig. 1C**) and EECs (**Fig. 1D**) from each patient, with similar plots for specific *GJ* genes shown in **Fig. S2** [top two rows of A (ESCs) and B (EECs)]. A Duncan analysis of the single-cell distributions shows that ESCs from all late stage endometriosis patients had significantly lower expression than ESCs from normal subjects and early stage endometriosis patients. The notable exceptions to this pattern in ESCs were those taken at the mid-secretory menstrual phase when progesterone levels, known to regulate connexin expression in the endometrium (Grummer et.al., 1994), are high [subjects 171 (normal) and 169 (late endometriosis)]. EECs show the inverse pattern that was even detectable in early stage endometriosis (patient 164), In EECs, the pattern was not influenced by the stage of the menstrual cycle when samples were collected, although did seem to be affected in the one non-cycling patient on birth control (003).

Comparison of individual GJ gene expression (**Fig. S1**) showed that *GJA1* (Cx43) was expressed most highly in both ESCs and EECs, almost 10 fold that of other connexins. For ESCs, all *GJ* genes measured, except *GJC2*, showed decreases in late stage endometriosis compared to normal subjects and early stage patients (**Fig. S1A**). In EECs, most of the *GJ* genes show the inverse behavior seen in ESCs, increasing expression in endometriosis compared to that seen in normal subjects. The exceptions were GJA1, the most highly expressed connexin, *GJA8, GJB7* and *GJC2* (**Fig. S1B**).

#### (b) Other genes

Several other genes involved in intercellular or cell-matrix interactions were also found to change in similar pattern to that of the GJ genes. The adhesion related genes *EpCAM*, ß-catenin (*CTNNB1*), as well as *NOV1* show decreased expression in ESCs in endometriosis (**Fig. 1E**) but increase in EECs (**Fig. 1F**), while the master transcriptional regulator of epithelial to mesenchymal transitions (EMT), Snail 1 (*SNAIl1*), which negatively regulates several adhesion molecules (Hugo et.al., 2011) and Cx43 (deBoer et.al., 2007), shows the inverse pattern (increasing with endometriosis in ESCs and decreasing in EECs). Several other genes change in one or other cells type. In ESCs, the metallo-endopeptidase, *CD10* (**Fig. S2A**), *NOV1* and the catalytic subunit of protein kinase A gamma (*PRKACG*) decrease, and ZO2 (*TJP2*), an accessory protein of tight and gap junctions, increases with disease (**Fig. 1E**). In EECs, only MAPK1 showed consistent increase in endometriosis patients (**Fig. 1F**). Some other genes showed increases in the two endometriosis patients in the proliferative or early secretory stages of their menstrual cycle [N-cadherin (*CDH2*), (**Fig. 1F**), and *PRKACG* (**Fig. S2B**)), but markedly different patterns in the non-cycling patient (**Fig. S2B**). Independent of these overall patterns, It is notable in the heat maps that the same subset of ESCs from the normal and early endometriosis samples (left three maps in **Fig. 1A**), and EECs from endometriosis patients (right two maps in **Fig. 1B**), that show high *GJ* genes expression, exhibit low expression of several of the tight junction associated genes, but higher expression of adhesion related genes (upper and lower rows, respectively in the group of genes labeled Adh/TJs) (**Fig. 1A and B**). Similar to what was observed for the tight junction associated genes, several of the regulatory genes (mostly kinases) we tested show the lowest expression in the cells that exhibit the highest expression of *GJ* genes (left three maps of ESCs (**Fig. 1A**) and right two maps if EECs (**Fig. 1B**).

### Cell cluster analysis

As an independent means to assess changes in gene expression in endometriosis, intra-sample cellular heterogeneity was analyzed by a graph-based cluster discovery algorithm (Ruan, 2009). By design, the algorithm is able to identify topologically distinct clusters and, importantly, can automatically determine the most appropriate number of clusters for each dataset. Utilizing this algorithm, ESCs and EECs were divided into 6 and 5 sub-populations, respectively (**Fig. 2A and C**). This analysis confirmed the general impression from the Heat maps, that the cell sub-groups characterized by high *GJ* gene expression (sc-6 for ESCs, and ec-5 for EECs) consistently decreased as a percentage of the ESC population (**Fig. 2B**), but increased as a percentage of the EEC population (**Fig. 2D**) from normal through early stage to late stage endometriosis. This cell cluster most reliably tracked the progression of endometriosis. Stated differently, the abundance of cells showing combined *GJ* expression levels at the 80^th^ percentile or above are highly predictive of disease state. For example, in ESCs all control subjects show >20%, early endometriosis 5-8% and late stage endometriosis <4%. This pattern was reversed in EECs, although we will need more patient samples to establish specific ranges.

**Figure 2:**
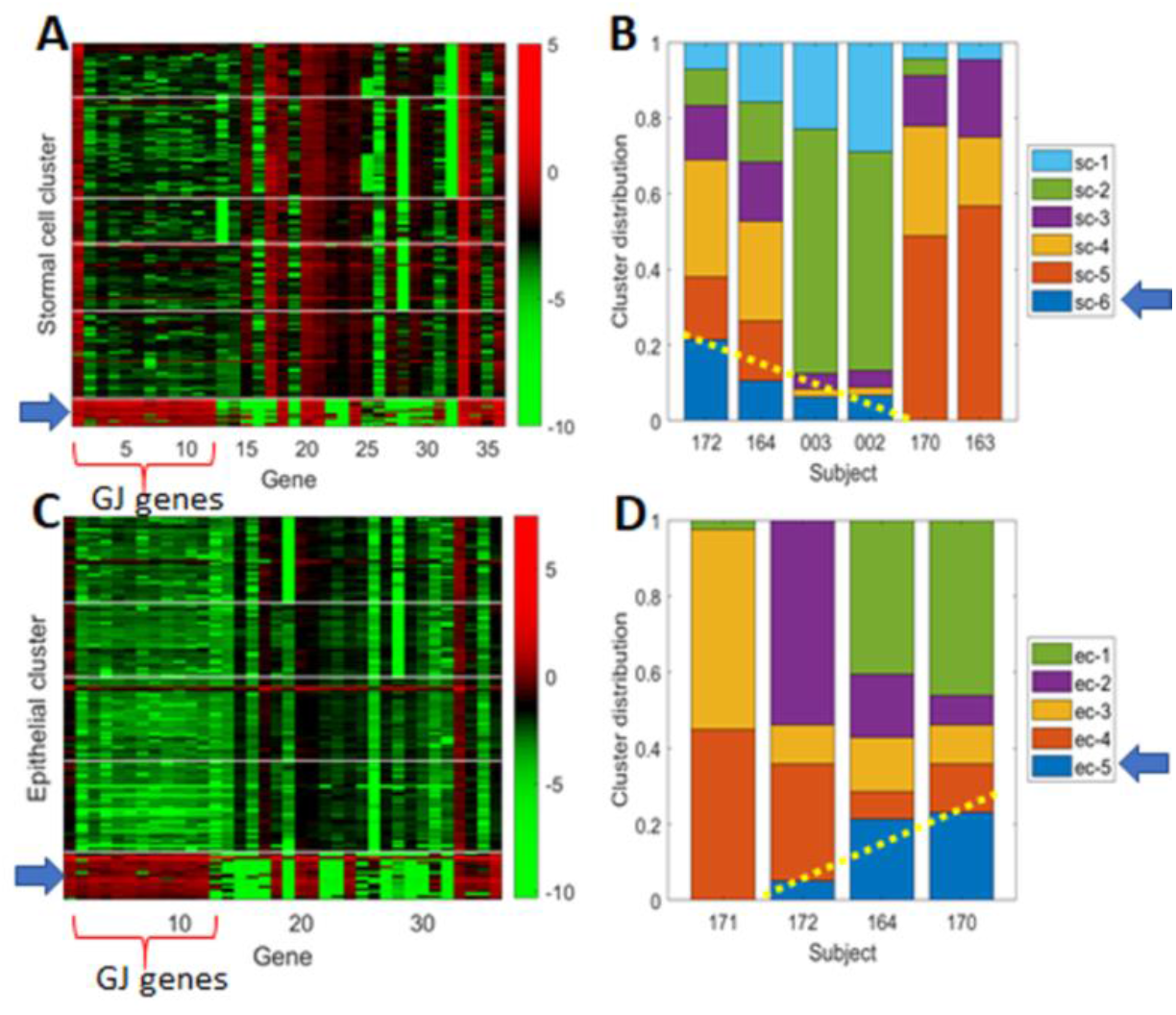
Cell clustering analysis comparing control and endometriosis samples show progressive changes in the fraction of cells characterized by high *GJ* expression. Cells from ESCs (**A**) and EECs (**C**) were grouped into 6 and 5 clusters, respectively, based on patterns of gene expression across all patients, with the bottom clusters [sc6 for ESCs, and ec5 for EECs] characterized by high GJ expression. Following the distribution of each cluster in all patients, endometriosis patients have consistently lower percentages of subpopulation sc-6 (**B**), and higher percentages of subpopulation ec-5 (**D**) compared to controls (trends highlighted by yellow dotted lines).

### Functional assessment of GJIC in ESCs and EECs, with endometriosis progression

We next assessed the functional consequences of these changes in *GJ* gene expression by measuring GJIC using an automated variant of the “parachute” technique where dye (calcein) loaded donor cells (D) are dropped onto a monolayer of acceptor cells (A) and the degree of dye spread to the monolayer (A/D ratio) is measured over 10-15 fields in an automated confocal microscope over time (**Fig. 3 A and B**). Given the changes in *GJ* gene expression between normal and endometriosis patients in **Fig. 1**, it was surprising that GJIC did not change significantly in either cell type, although a small decrease was seen in ESCs and increase in EECs (**Fig. 3C**). However, as endometriosis is characterized by invasive behaviors following interactions with the peritoneal mesothelium, we also assessed heterotypic GJIC between endometrial cells and a peritoneal mesothelial cell (PMC) line (LP9). This revealed a dramatic induction of GJIC that was most notable in ESCs from endometriosis patients (4 fold increase compared to 2 fold in normal subjects). While some induction was also observed in EECs, this was not significant in either normal or endometriosis samples (**Fig. 3C**). This induction occurred relatively rapidly, within the context of the assay (~ 2 hours).

**Figure 3:**
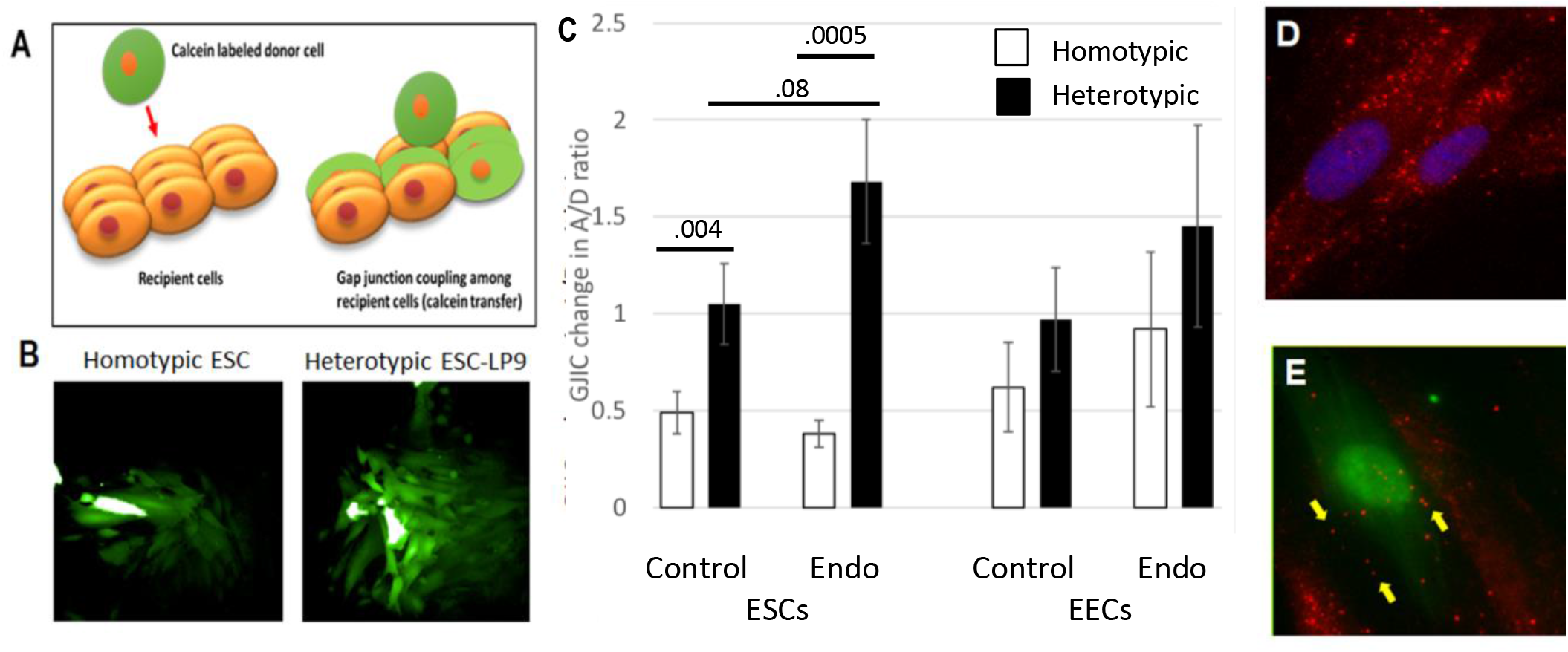
Heterotypic coupling between ESCs and PMCs is dramatically induced in endometriosis. Gap junction intercellular coupling (GJIC) was measured by a modified “parachute assay” where calcein ester loaded donors are dropped onto a monolayer of acceptors, and dye transfer measured over time as a linear increase in fluorescent acceptor/donor ratio (**A**). This can measure homotypic (donor and acceptor the same), or heterotypic (endometrial donors and mesothelial acceptors) GJIC (**B**). Although there are slight changes in homotypic GJIC in ESCs and EECs consistent with the changes in expression levels of connexins, they were not significant (**C**-green bars). By contrast, heterotypic interactions with mesothelial cells (PMCs) caused a significant induction of GJIC, most notably in ESCs from endometriosis (**C**-black bars). Immunocytochemistry showed this was likely due to a redistribution of Cx43 (red) from primarily cytoplasmic in homotypic ESCs (**D**), to the interfaces of ESCs (green) and PMCs in heterotypic cultures (**E**). Samples tested: ESC control (8) and endometriosis (11); EEC control (8) and endometriosis (5), which included 2-4 biological replicates for each patient. Significance level was calculated by two tailed t-test.

Given this time course, which seemed inconsistent with a transcriptional event, we examined the distribution of Cx43 protein in ESCs alone, and after contact with PMCs. Alone, both stromal (**Fig. 3D**) and mesothelial cells (not shown) show primarily intracellular Cx43 staining. By contrast, in heterotypic cultures, very little intracellular staining is seen within ESCs (fluorescent green labelled cell in **Fig. 3E**), but plaques are readily observed at sites of contact with PMCs (yellow arrows, **Fig. 3E**), although significant intracellular labeling within PMCs was still evident. This suggests that the heterotypic contact may trigger enhanced trafficking of Cx43 to the cell surface.

Since the major functional difference we observed related to gap junctions in samples from endometriosis patients were associated with heterotypic contacts with mesothelial cells, especially by ESCs, we predicted that they may influence differential adhesion between the cells. To measure the degree to which heterotypic adhesion properties with mesothelial cells differ between EECs and ESCs, we used atomic force microscopy (AFM) that enables quantitative assessment of the extent of intercellular interactions (Sancho et. al., 2017; Roca-Cusachs et.al., 2017). Adhesiveness between PMCs (LP9 cells), and EECs and ESCs from endometriosis patients was measured by attaching a single cell to a cantilever of an AFM probe (**Fig. 4A**). The attached cell was placed in contact with a monolayer of either the same cell, or a different cell type, growing on a culture dish, and the force needed to separate the probe-attached cell from the plate-attached cell was measured (**Fig. 4B**). Mesothelial cells show low levels of adhesion to one another that was similar to their adhesion to EECs. In contrast, ESCs showed significantly higher adhesion to PMCs (**Fig. 4C**). This data is consistent with ESCs being the cell type that is the major invasive front into the mesothelium (Nair et.al., 2008; Burney and Giudice, 2012).

**Figure 4:**
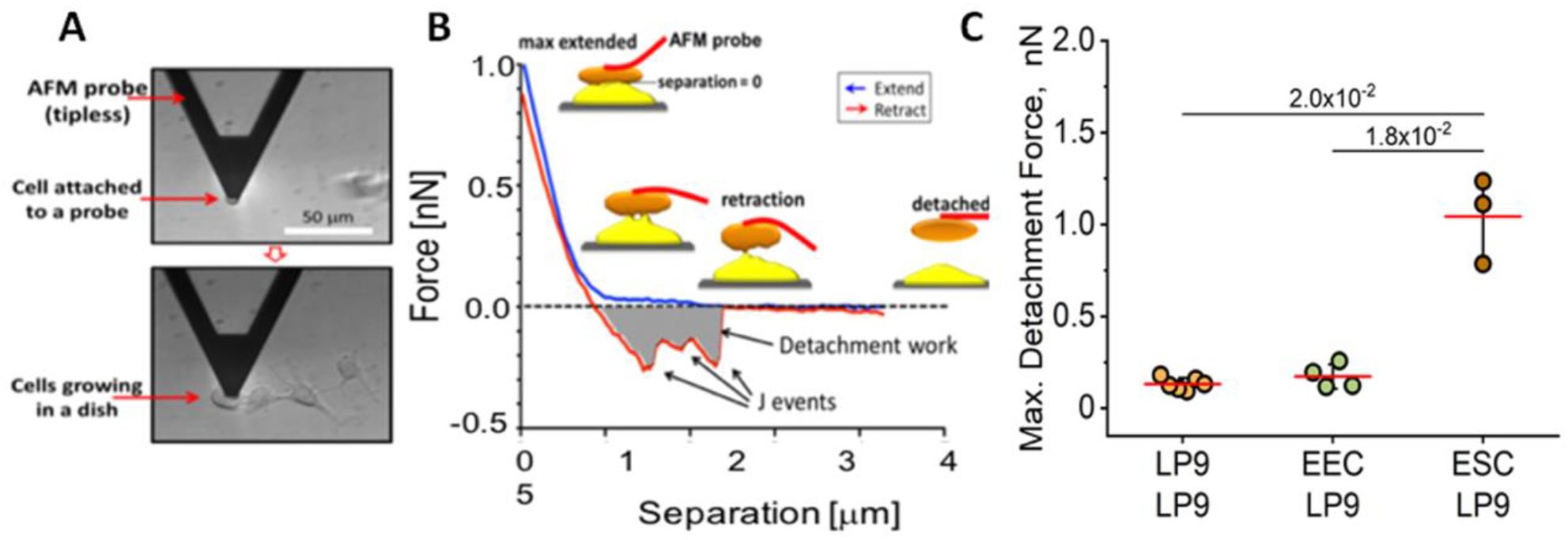
Atomic Force Microcopy (AFM) demonstrates enhanced adhesion between ESCs and PMCs. Cells attached to an AFM tip are bought into contact with PMCs growing on a dish (**A**), and the force needed to separate them is measured (**B**). PMCs (LP9 cells) show similar adhesion to one another as to EECs, but much stronger adhesion to ESCs (**C**). Significance determined by two tailed t-test

### Invasive Potential of Endometrial Cells from Normal Subjects and Endometriosis Patients, and dependence on GJIC

We then directly assessed the invasiveness of primary endometrial cells from normal and endometriosis patients using an established 3D-invasion assay that assessed the efficiency with which ESCs or EECs can pass through a confluent monolayer of PMCs grown on a growth-hormone depleted Matrigel-coated membrane (**Fig. 5A**). Initial studies using unseparated endometrial cells showed significantly greater invasiveness in samples from endometrial patients than control subjects (n=3; **Fig. 5B**). For ESCs (n=3 and 6 for control and endometriosis, respectively) and EECs (n=3 and 6 for normal and endometriosis, respectively) the differences were not significant. In all cases tested the level of invasion was not seen when endometrial cells were dropped on a membrane alone, so that the invasion was dependent on contact with a mesothelial monolayer (data not shown).

**Figure 5:**
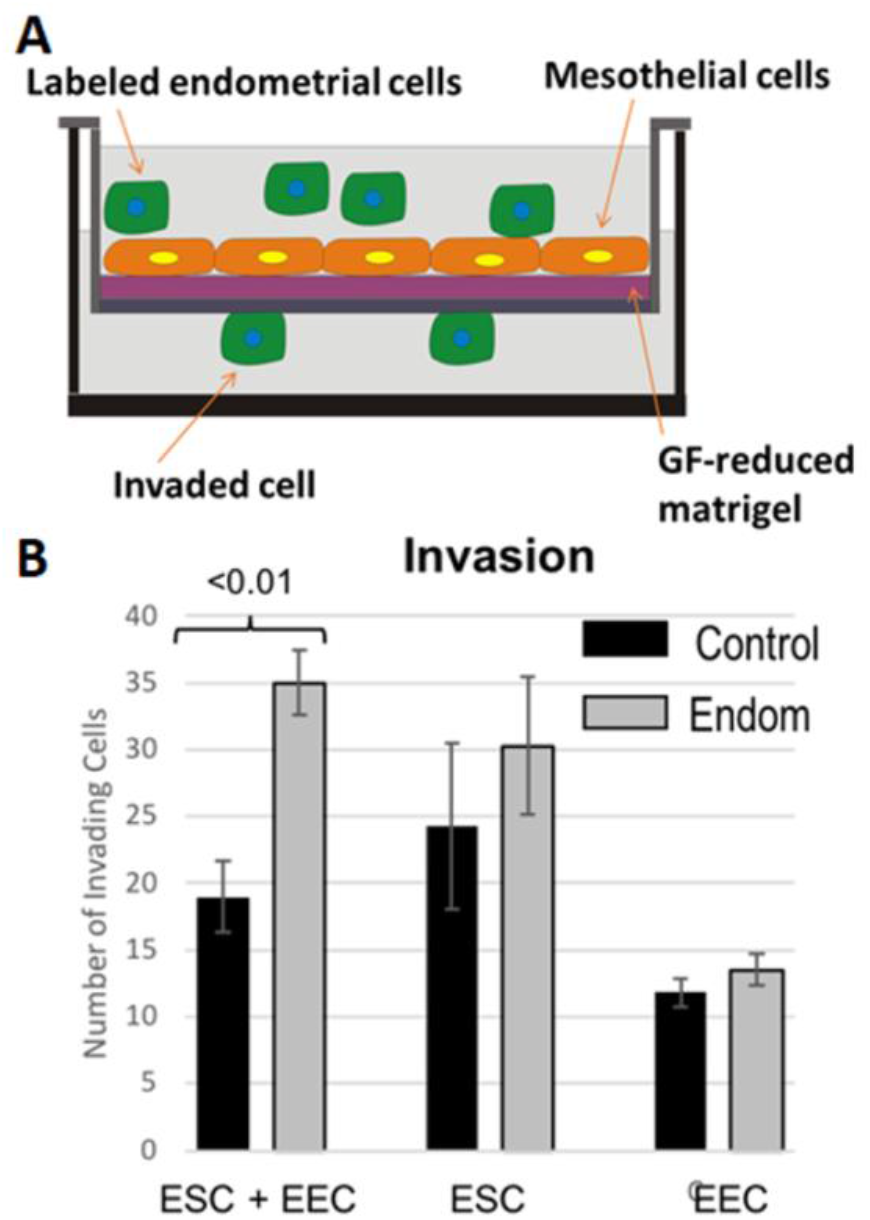
Enhanced invasiveness of endometrial cells from endometriosis patients across a mesothelial monolayer. Using a 3-D model of endometrial cell invasion across a peritoneal cell monolayer (**A**), endometrial cells from patients show greater invasion than those from control subjects that was significant with a mix of ESCs and EECs (**B**). Number of independent patients tested - All Controls (3); Endometriosis – ESC+EEC (3 patients); ESC (5); EEC (4). Significance was determined by twotailed t-test.

Given that we had demonstrated that we had demonstrated that endometrial cell contact with mesothelial cells specifically induced GJIC, we tested the degree to which invasive behavior was dependent on GJIC by targeting Cx43, the dominantly expressed Cx in both cell types (**Fig. S2**). Four approaches were used to test this. Firstly, we pre-treated both PMCs and endometrial cells (not separated into ESCs and EECs) with GAP27, a peptidomimetic to the extracellular domain of Cx43 that we, and many others (Evans and Leybaert, 2007), showed blocked the formation of gap junctions between newly contacting cells, in this case by ~85%. This reduced invasiveness of unseparated endometrial cells from control subjects by 57% (although not significant at the 0.05 level), and from endometriosis patients by 70% (p<0.01) (**Fig. 6A**).

**Figure 6:**
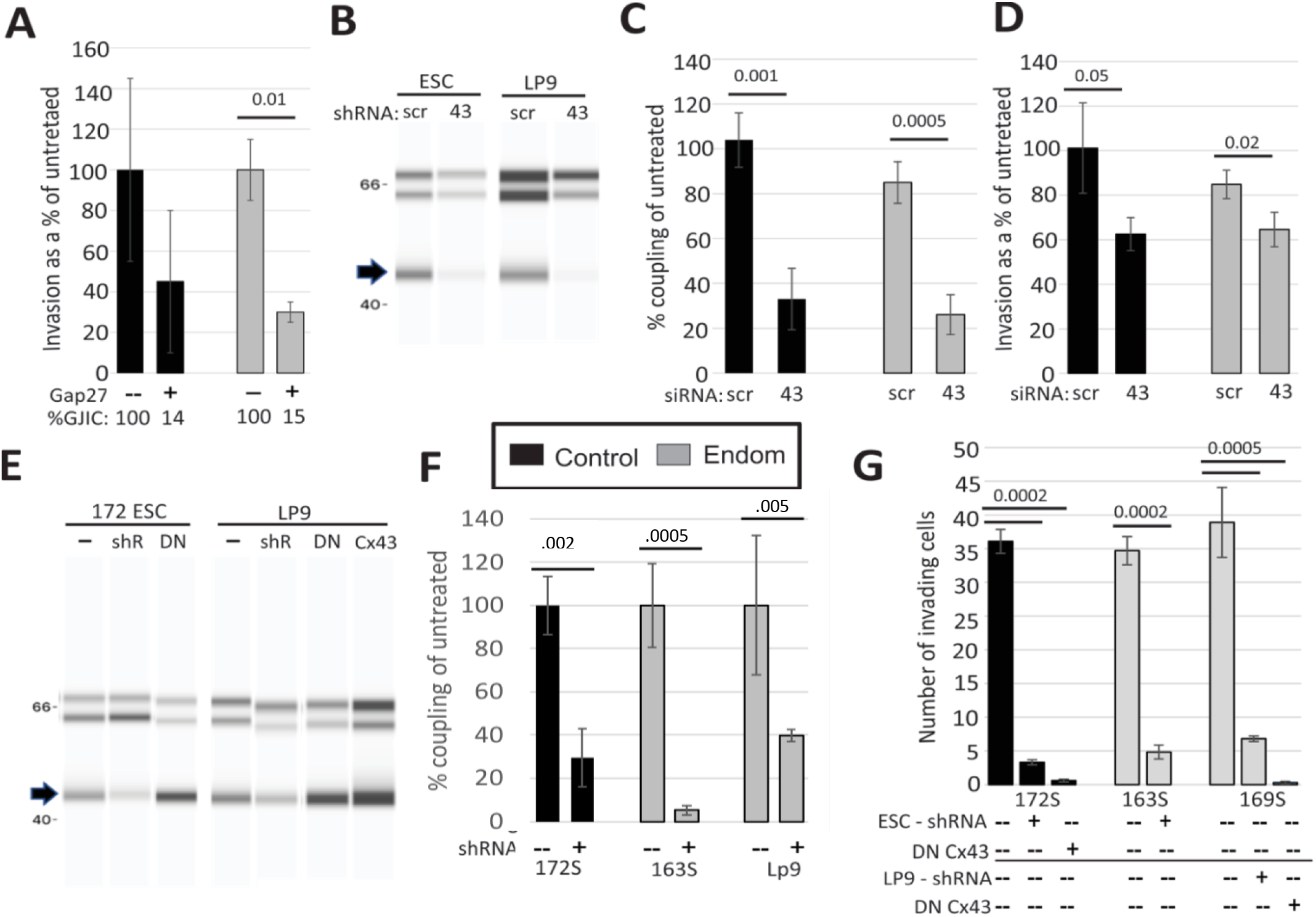
Invasiveness of ESCs across a PMC monolayer is dependent on Cx43 GJIC: (**A**) Averaging ESCs from all control (black bars, n=3) and endometriosis patients (grey bars, n=6), invasion was inhibited by a peptide inhibitor of GJ channels, GAP27 (percent GJIC compared to untreated shown below each bar). (**B - D**) Cx43 siRNA transfection of either ESCs or PMCs (LP9) reduced protein levels of Cx43 (arrow) compared to Laminin A and B controls (upper bands) (**B**), inhibited heterotypic GJIC compared to scrambled siRNA [Control n=5 (independent replicates from 2 patients); Endo n=8 (independent replicates from 5 patients)] (**C**), and also inhibited invasion [Control n=6 (2-3 independent replicates of two patients); Endo n=14 (2-4 independent replicates of 6 patients)] (**D**). (**E – G**) Stable, doxycycline inducible shRNA infections of ESCs from a normal (172) and endometriosis (163) patient, as well as PMCs (LP9), reduced levels of Cx43 (arrow) compared to Laminin controls (**E**), inhibited GJIC (n=3-7 independent experiments with 3 different shRNAs) (**F**) and invasiveness by ~90% [10 technical replicates of all samples; 3-4 biological repeats for +/-shRNA using 3 different shRNAs (controls were internal)] (**G**). The block of invasiveness was observed independent of whether Cx43 expression was inhibited in ESCs or PMCs. DN Cx43 (which blocks channel function, but not protein assembly into GJs) was also expressed in ESCs and PMCs at approximately equal levels to wt Cx43 (total Cx43 levels doubled - **E**). This caused an even greater inhibition of invasiveness (~98% - **G**). All statistical significance based on onetailed t-tests.

Secondly, we tested purified ESCs (as only they showed increased adhesiveness to PMCs (**Fig. 4**)), using a combination of two siRNAs targeted to Cx43, where the PMC monolayer was transfected immediately prior to the invasion assay. This caused an average of 65 ± 9 % reduction in Cx43 protein (**Fig. 6B**) and ~70% reduction in GJIC between endometrial and mesothelial cells (p<0.001 in control and <0.0005 in endometriosis) (**Fig. 6C**). Invasiveness was reduced by 30-40% (p<0.05 in control, and <0.02 in endometrial samples) (**Fig. 6D**). In all cases, Cx43 siRNA effects were compared to effects by either a scrambled siRNA, or siRNA directed to GAPDH.

Thirdly, in order to avoid any negative effects of the transient transfection process for siRNA on the integrity of the PMC monolayer, or invasive potential of ESCs, we prepared stably expressing LP9 cells and ESCs from a normal and endometriosis patient (172 and 163, respectively), by infection with lentivirus that either expressed shRNAs targeted to Cx43 that were inducible by doxycycline. In the presence of doxycycline, shRNA suppressed Cx43 protein levels by 35 ± 6 % (**Fig. 6E**), and GJIC was by 88 ± 4 % in the cell types tested (**Fig. 6F**). Invasive behavior was also inhibited by 86 ± 2 % when Cx43 was suppressed in the ESCs from either control or endometriosis patients, or in the PMCs (**Fig. 6G**).

Finally, we used the same Lentivirus system to express a dominant negative Cx43 construct, Cx43 T154A (DN Cx43), which we have previously shown preserves normal gap junctional plaque structures, but fails to open functional channels, and prevents the opening of co-expressed wt Cx43 (Beahm et.al, 2006). Expression of DN Cx43 increased total Cx43 levels by 2 fold in ESCs and 1.4 fold in PMCs (**Fig. 6E**), decreased GJIC by 90% (**Fig. 6F**) and invasive behavior by a remarkable 98-99%, whether expressed in ESCs or PMCs (**Fig. 6G**).

This analysis clearly shows that both EECs and ESCs can be invasive across a mesothelium, and that this is even true when they are derived from control subjects, although the levels are higher in cells from endometriosis patients. Three independent treatments that specifically block either functional GJIC between ESCs and PMCs (GAP27 or DN CX43) or expression of Cx43 in either of the cell types, greatly reduce the invasive behavior in both control and endometrial samples. Tests on EEC invasiveness (data not shown) showed similar dependence of invasiveness on Cx43, but the higher variability between samples from different patients led to the effect not being significant at the 0.05 level.

### AFM analysis of ESC effects on the PMC monolayer

Given the clear dependence of invasiveness of ESCs through the mesothelium on Cx43 mediated GJIC, we turned to AFM to provide insights into how this occurs. We specifically probed the influence of ESCs, in the presence or absence of Cx43, on the “barrier function” of the mesothelium (i.e. the “tightness” of contact between PMCs that prevent transmigration of cells). ESCs from a control subject (172) or endometriosis patients (169 and 170) were first labeled with the membrane dye DiO, and dropped onto a PMC monolayer at a ratio (ESC:PMC) of 1:20. After ~3 hrs, the monolayer was imaged with AFM by pressing it with a ‘sharp’ conical probe at a constant pressure of 1 nN to obtain a 3-D contour map of the monolayer (**Figs. 7A-B**). This readily allowed the interfaces between cells to be identified (**Fig. 7C**), and to measure the depth of penetrance between cells. ESCs from several patients all induced an increase in penetrance, measured ~10um away (~2 cell diameters) from site of the dropped cells, identified based on prior DiO labeling (**Fig. 7D**). This was also evident as a widening in the gap between cells (compare **Figs 7A and B**). This effect on penetrance between cells was most pronounced in ESCs from an endometriosis patient (169) compared to a control patient (171) (**Fig. 7D**), and also showed a dependence on the density of ESCs, as penetrance was less when a 1:50 ratio of ESCs:PMCs was used instead of 1:20 (data not shown). We have also conducted a similar test using an AFM probe to which a 3 μM diameter glass bead is attached to mimic the shape of a cell, and similarly shown that ESCs induce greater ability for larger objects to penetrate the monolayer (data not shown).

**Figure 7:**
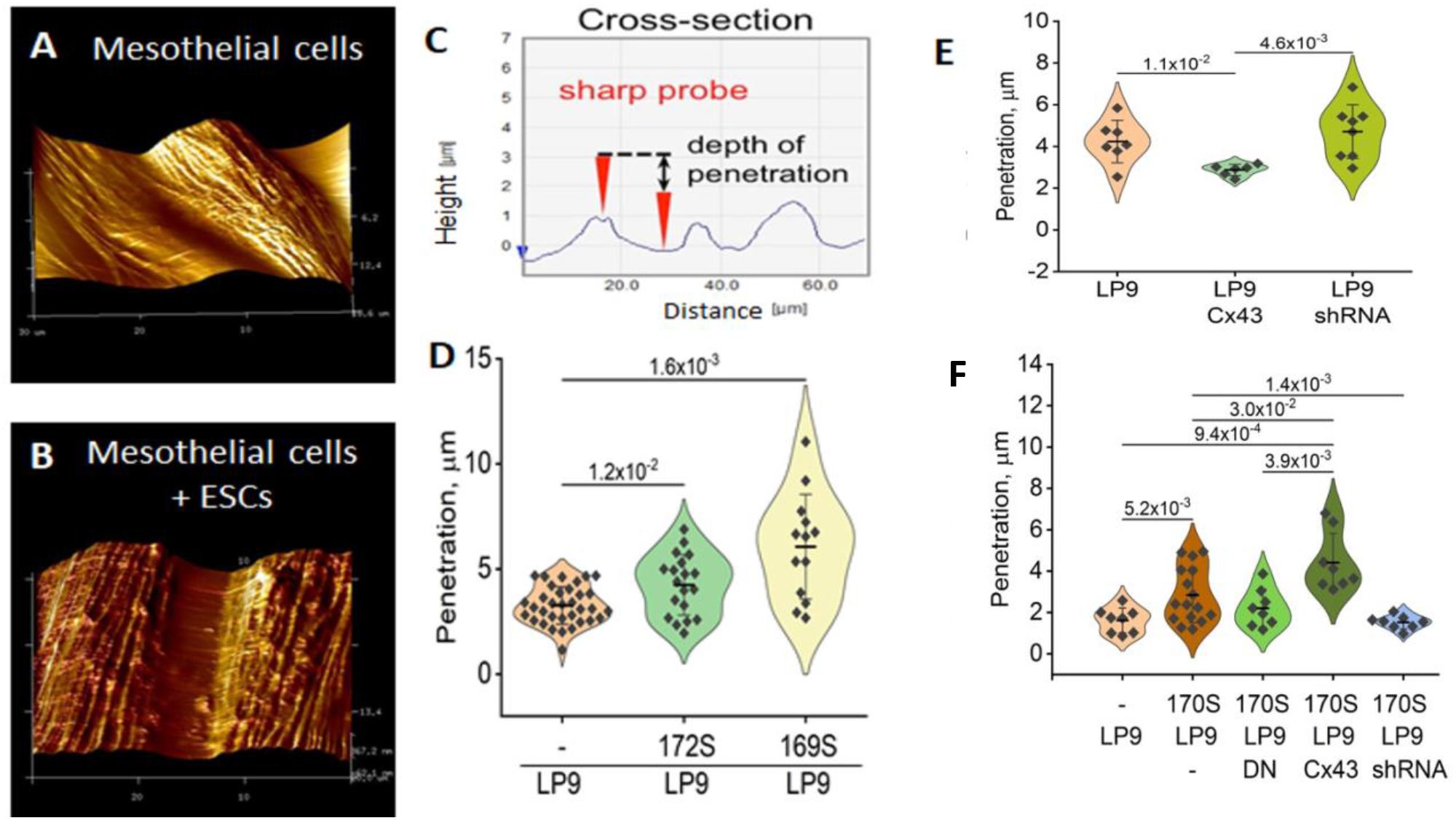
AFM demonstrates that ESCs induce disruption of the barrier function of a mesothelial monolayer, which is dependent on Cx43 channels. The topology of the mesothelial monolayer surface shows a significant increase in the spacing between cells in the presence of ESCs (**A-B**). Using constant pressure, the tip can measure the depth of penetration between cells (**C**). This was increased from that measured in anLP9 PMC monolayer alone (n=25) when ESCs were dropped onto the monolayer, particularly from an endometriosis patient (169, n=19)) compared to control (172, n=13) (**D**). Cx43 appears to help preserve the barrier function of the mesothelium alone, as overexpression reduced penetration, while Cx43 shRNA infection increased penetration in the PMC monolayer in the absence of ESCs (**E**). By contrast, in the presence of ESCs from a patient (170), Cx43 overexpression in PMCs enhances penetration, while blocking Cx43 with either shRNA or DN Cx43 neutralize the effect of ESCs (**F**). Each dot represents a cell measurement from plates prepared on the same day for comparison. Statistical tests of significance by two-tailed t-test (see methods).

We then used the shRNA, DN Cx43 and wt Cx43 infected LP9 cells, characterized in **Fig. 6E**, to test the dependence of these changes on Cx43 GJIC. First, we observed that the “barrier function” of mesothelial cells alone (i.e. in the absence of ESCs) was dependent on Cx43 expression, as the degree of penetrance was reduced when Cx43 was overexpressed, and increased when Cx43 was inhibited by shRNA (**Fig. 7E**). Strikingly, this effect was exactly inverted when we introduced ESCs, as now Cx43 overexpression significantly increased penetrance, while Cx43 inhibition by shRNA completely eliminated the effect of ESCs, so that penetrance was the same as in their absence. This effect was due to the channel forming role of Cx43, as the same reduction in penetrance was observed in LP9 cells infected with a DN Cx43, which would preserve, or even enhance, any adhesive functions of Cx43 between ESCs and PMCs.

## DISCUSSION

Invasive processes of any cell into a “foreign” environment occurs in many instances, both normal (during development, extravasation, blastocyst implantation) and pathological (metastasis) and involve close intercellular interactions. While the need for initial adhesion events has long been recognized, it has become increasingly evident that the formation of heterotypic gap junctions between the invading and target tissue is an early event in implantation (Grummer et.al., 1996), extravasation (el Sabban and Pauli, 1991, 1994; Ito et.al, 2000, Naoi et.al, 2007) and metastasis (Lamiche et.al., 2012; Stoletov, et.al., 2013; Hong et.al, 2015; Cheng et.al., 2015). However, despite endometriosis being the most common form of pathological invasiveness (affecting 10% of women worldwide), there are few studies on the molecular characterization of gap junctions, (or other intercellular junction components) in this disease. Herein, we sought to determine if dysregulated expression of these endometrial genes associated with disease status

### Expression of GJ and other genes

In a test sample of 6 endometriosis patients and 2 healthy controls, we found highly significant shifts in the expression of several junctional genes and proteins that regulate junctional contacts transcriptionally and post-transcriptionally. The most predictive change associated with endometriosis was in the gap junction (*GJ*) gene family encoding connexins, specifically in terms of the subpopulation of cells showing high *GJ* expression (**Fig. 1A-D**). This became even more evident in a cell clustering analysis, which demonstrated that the only group of cells where fraction of the cell population reliably tracked disease progression were those characterized by high *GJ* expression with (**Fig. 2**). This pattern has likely escaped prior screens of endometrial tissue using bulk PCR approaches, as the changes in *GJ* gene expression in the two major cell types of the endometrium were in opposite directions, with stromal cells showing decreased expression (**Fig 1C**), and epithelial cells evidencing upregulation (**Fig 1D**) as disease stage increased.

Notably, these changes were not restricted to the connexins expressed most abundantly in the endometrium (Cx43, 26 and 32), but applied to most of the *GJ* gene members screened. It was also notable that Cx43 was the dominantly expressed Cx in both ESCs and EECs. For the former, this was consistent with what has been reported in the literature, but EECs have been reported to display mostly Cx26 and Cx32 expression in eutopic endometrium (Jahn et.al., 1995). This has been shown to switch to Cx43 in ectopic lesions (Regidor et.al., 1997), and even eutopically in a baboon model of endometriosis (Winterhager et.al., 2009). While it is possible that the short time in culture modified this expression pattern of EECs in our study, it is also possible that the switch to Cx43 occurs very early in release of cells from the endometrial lining, even in normal subjects. In either event it is clear that the expression of Cx43 is greatly enhanced in EECs from endometriosis patients compared to control, even while they are still resident in the eutopic endometrium.

In terms of other genes involved in intercellular interactions, many do not show consistent changes, at least in both cell types, but two genes associated with adhesion (EpCAM and ß-catenin) show the same expression shifts as the *GJ* genes, while Snail 1, a master transcriptional repressor that promotes endothelial to mesothelial transition (EMT), shows the opposite pattern. This is consistent with the demonstrated suppressive effect Snail 1 has been shown to have on Cx43 (deBoer et.al., 2007), and the promotion of a migratory phenotype in ESCs, and its repression in EECs. However, it is not clear if in this instance Snail1 is activating a typical EMT response, as the expression of E-cadherin, which is a primary target for suppression by Snail1, was not detected in the endometrium, and N-cadherin, the expression of which is normally increased by Snail1, increases in endometriosis EECs coincident with a Snail 1 decrease. Furthermore, the increase in Snail1 in ESCs is associated with a reduction in ß-catenin levels, which may limit EMT effects as they typically depend on activation of ß-catenin signaling. This is consistent with the view that EMT transitions in endometriosis are only partial (Konrad et.al., 2020).

With regard to kinases that regulate many intercellular junctional components, it is notable that the cells with the highest *GJ* expression (control ESCs and endometriosis EECs) in general show the lowest expression of most kinase genes (see heat maps in **Fig. 1 A-B**), suggesting an inhibitory relationship. However, in terms of global kinase levels, the only kinases to show significant shifts with endometriosis were PKA gamma in both ESCs and EECs, and MAPK1 in EECs (**Fig. 1E-F, and Fig. S2B**), all of which changed in the same direction as the GJ genes. This is a striking example of how overall expression levels can be misleading when compared to the details of expression patterns at the single cell level.

### GJIC and invasiveness

These expression profiles initially pose an interesting conundrum, as stromal cells are thought to be the primary compartment involved in endometriotic lesion formation (Lucidi et.al., 2005; Burney and Giudice, 2012), and show the greatest EMT-like response in endometriosis (see above), yet they also show repressed levels of Cx43, which is needed for invasive behavior in metastasis and extravasation. However, functional studies reveal that ESCs, while poorly coupled homotypically, show a dramatic induction of heterotypic coupling when contacting mesothelial cells that is greatly enhanced in endometriosis samples (**Fig. 3C**). This induction, linked to increased trafficking of Cx43 to the cell surface (**Fig. 3D and E**), is very analogous to what has been observed in breast cancer, where internalized Cx43 and 26 in the primary tumor, traffics to the surface in lymph node metastases (Kanczuga-Koda et.al., 2006). Enhanced heterotypic coupling with capillary endothelia (el Sabban and Pauli, 1991, 1994; Ito et.al., 2000) or target tissues (Lamiche et.al., 2012; Hong et.al., 2015) have been shown to be critical to invasive behavior during early metastasis.

The importance of the enhanced heterotypic coupling between ESCs and PMCs, unique to endometriosis, to their invasiveness across the mesothelium was demonstrated using four different modes of inhibition of Cx43. Specifically, invasion was not only inhibited by suppression of Cx43 expression (**Fig. 6D and G**), but also by a peptide blocker GAP27 (**Fig. 6A**), which inhibits all channel function, but leaves protein expression unaffected, and by DN Cx43 expression (**Fig. 6G**), which actually increases protein levels (**Fig. 6E**) and gap junction structures (Beahm et.al., 2006), but blocks all channel activity. Notably, inhibition of invasion is the same whether Cx43 function is ablated in ESCs or PMCs (cf. shRNA and DN Cx43 treatments for 163S and 169S in **Fig. 6G**). This suggests that heterotypic GJIC likely triggers the invasive behavior, which, as we demonstrate with AFM (**Fig. 7**), is associated with increased separation between PMCs, likely resulting from a disruption of the adhesion and tight junctions between the cells (i.e. breakdown of the “barrier function”), akin to what happens during extravasation (Ito et.al., 2000; Reymond et.al., 2013).

In **Fig. 8** we present a summary of how interactions between the mesothelium and endometrial cells arriving in the peritoneum by retrograde menstruation differ in normal and endometriosis patients, resulting in lesion formation. The enhanced GJIC seen in endometriosis ESCs (**Fig. 8B**) would allow signals, indicated by green triangles, to pass effectively from endometrial cells to the mesothelium, where they are further propagated through GJIC, to promote disruption of “barrier function” between PMCs, allowing invasion. Interestingly, in the absence of signals from ESCs, GJIC was shown to be important to maintenance of the barrier function (**Fig. 7E**). This may also be why DN Cx43 was more effective in preventing invasion than Cx43 knock-down by sh- or si-RNA (**Fig. 6G**), as it may reinforce the barrier functions by enhancing adhesiveness between PMCs. The model emphasizes how increased receptiveness of the ESCs to form heterocellular gap junctions in endometriosis can lead to invasive lesion formation. However, it is also possible that they endometriotic ESCs may make more of the hypothetical signals that induce disruption of the barrier function, emphasizing the importance in identifying such factors. It should be noted that we cannot definitively exclude involvement of factors released through Cx43 hemichannels in the invasive process, but there are no obvious signals that would induce their opening in these experiments, and effects on invasion are the same independent of the cell in which Cx43 is inhibited.

**Figure 8:**
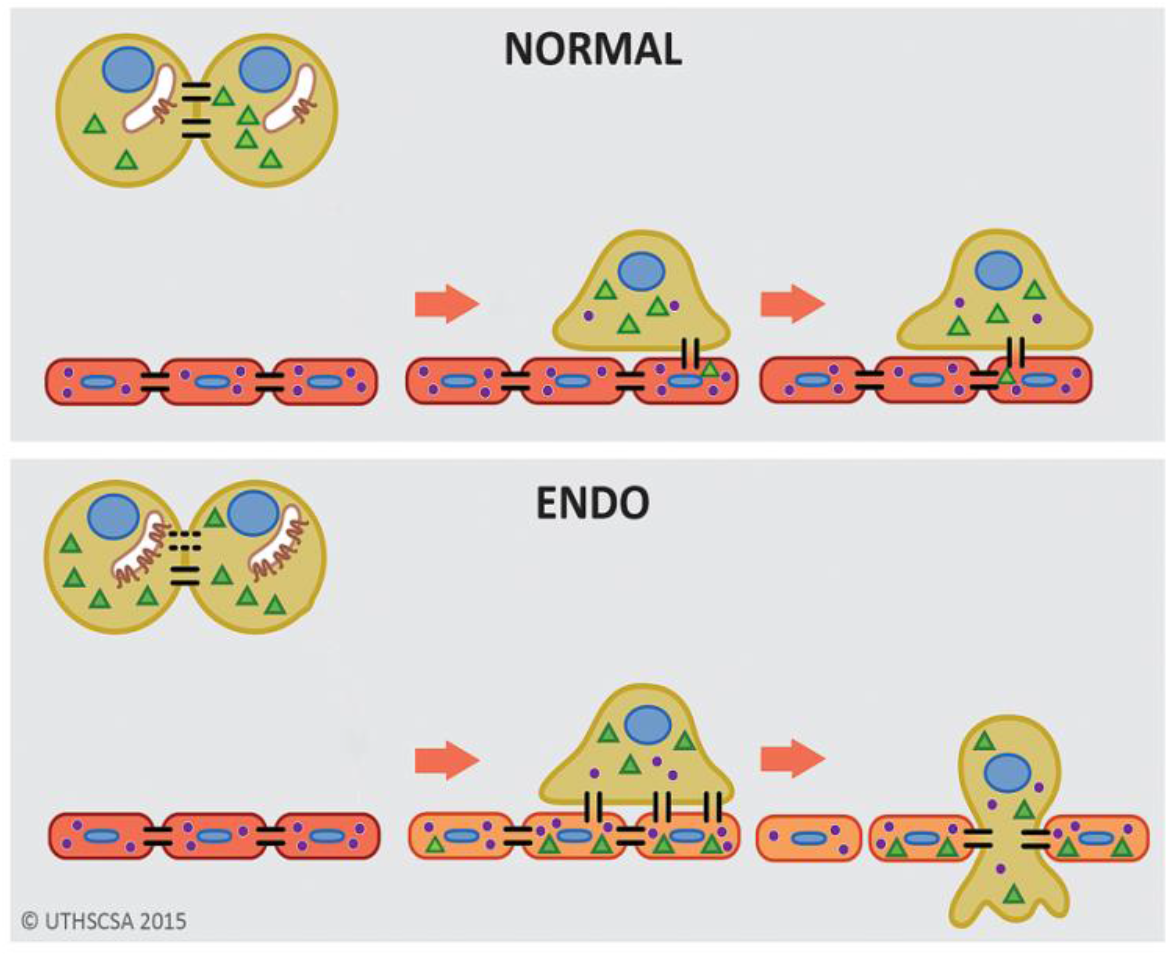
Model of GJIC induction of trans-mesothelial invasion. (**A**) In healthy patients, when endometrial cells (brown) encounter a mesothelium (following arrival in the peritoneum via retrograde menstruation), modest GJIC provides limited exchange of signals from ESCs to PMCs (green triangles) or PMCs to ESCs (purple dots). (**B**) In endometriosis, endometrial cells have Cx43 mostly in intracellular stores, but encounters with mesothelial cells triggers Cx43 trafficking to the cell surface. The increased GJIC that results mediates transfer of signals to PMCs (green triangles), which propagate through the mesothelium, inducing disruption of the adhesive and tight junctions between PMCs, facilitating invasion of the ESCs.

Much remains to be done to identify the intercellular signals that mediate barrier function disruption, and also to understand reciprocal effects of the mesothelium on the ESCs, aside from the demonstrated induction of heterotypic gap junction formation. However, the current study clearly implicates changes in gap junction expression in the earliest phases of the development of endometriosis, and their critical role in initiating lesion formation. They also demonstrate that changes within the uterine endometrium prime these cells to be invasive once they reach the peritoneal cavity, in much the same way that the metastatic potential of cancer cells is determined as they leave the primary tumor, suggesting that understanding each of these processes will inform the other.

## MATERIALS AND METHODS

### Primary endometrial epithelial cell isolation from endometrial biopsies

Primary ESCs and EECs were isolated from endometrial biopsies obtained from women with and without endometriosis. All women provided informed consent prior to participating in this Institutional Review Board approved protocol. Study subjects were premenopausal women between 20 and 45 years of age with regular menstrual cycles undergoing laparoscopic surgery for gynecologic indications (**Table 1**). Women with pelvic inflammatory disease/hydrosalpinx, endometrial polyps, or submucosal fibroids were excluded. Endometriosis was staged according to the revised ASRM criteria and confirmed by histopathologic review of peritoneal or cyst wall biopsy in all cases. Fertile women undergoing tubal sterilization and without endometriosis at surgery were considered healthy controls. Menstrual cycle phase (proliferative or secretory) was determined by cycle history and confirmed by serum estradiol and progesterone levels. Endometrial tissue was obtained by pipelle biopsy at the time of laparoscopic surgery.

The biopsy material was dissociated by shaking in 5mg/ml collagenase and 2.5mg/ml DNase in Hanks Balanced Salt Solution at 37^0^C for 1 hour. Isolation of primary ESCs and EECs from the biopsies was performed using a combination of straining (45uM nylon filter) and differential sedimentation (EECs cluster and sediment faster), followed by differential attachment (EECs adhere less well to culture plates), in a modification of the method developed by Kirk and Irwin (1980) used in prior studies De La Garza, et.al., 2012; Chen et.al., 2016). In some experiments the differential attachment step was replaced by using an Ep-CAM affinity column to enrich EECs. Both methods achieve about 97% purity for EECs and ESCs, as illustrated in **Fig. S1** by immunostaining for epithelial [EpCAM (ab71916 from Abcam) and CK 7 (ab902 and 1598 from Abcam)] and stromal [Vimentin (MA1-10459 from Thermo Fisher; NBP1-92687 from NovusBio)] markers. Gene analysis was performed exclusively on P0 or P1 cells, while functional tests of coupling and invasion used cells between P0 and P5 (ESCs) or P0 to P3 (EECs).

### Cell Culture

Primary ESCs were cultured in Dulbecco’s Modified Eagle Medium (DMEM)/F12 (1:1) (Gibco, Buffalo, NY) containing antibiotic/antimycotic mix (Gibco, Buffalo, NY), 10 μg/ml insulin (Sigma, St. Louis, MO) and 10% heat inactivated fetal bovine serum (FBS - Gibco, Buffalo, NY) as described previously (Ferreira *et al*., 2008). EECs were cultured in MCDB/Medium 199/MEMα (1:1:0.6) containing antibiotic/antimycotic mix, 10ug/ml insulin, D-Glucose (0.45%) (Sigma, St. Louis, MO), GlutaMax and 10% FBS (Gibco, NY). Prolonged culture was in defined KSFM with supplement, 1% FCS, and antibiotics/antimycotics (Gibco, NY) to preserve differentiated state of the EECs (Chen et.al., 2016) although this generally was only possible to 3 - 4 passages. All experiments were performed using low passages (≤5 for ESCs ≤3) to avoid loss of differentiated characteristics. Established LP9 cells (Corriell Cell Repositories, Camden, NJ) were used as a model for peritoneal mesothelial cells and cultured as described previously (De La Garza, 2012, Liu *et al*., 2009) and grown in MCDB 131.Medium 199 (1:1 - Gibco, NY) with 15% FBS, sodium pyruvate, GlutaMax, antibiotic/antimycotic mix (Gibco, NY), hEGF (20ng/ml) and hydrocortisone (0.4ng/ml) from Sigma, St. Louis, MO. Previous studies, including our work, have validated and used LP9 cells as a model for peritoneal invasion by endometrial cells (Nair et.al., 2008)

### Single-cell RNA expression of a connexin gene panel by microfluidic PCR

Automated single cell isolation and processing for cDNA amplification was performed in the C1 (Fluidigm Inc) system, using the C1 integrated fluidic circuit (IFC) chips. Captured single cells were visually confirmed by viewing the microfluidic wells of the C1 IFCs on an inverted light microscope. Empty wells or those containing doublets were excluded. Single-cell cDNA from the IFC chips is then subjected to microfluidic PCR amplification in the Biomark platform (Fluidigm, Inc), using Biomark PCR IFC chips with1936 (48×48) or 9216 (96×96) microfluidic wells (Polusani et.al., 2019). The gap junction gene panel and corresponding PCR primer sequences for expression profiling selected from the PrimerBank database (http://pga.mgh.harvard.edu/primerbank) are outlined in Supplemental Table 1. In each Biomark chip assay, universal RNA (200 pg) from human normal tissues (cat #4234565, BioChain, Newark, CA) and no template control (NTC) served as positive and negative controls, respectively. Valid PCR products were determined by amplicon melting temperature curves of each gene. Expression values were determined based on cycle threshold (Ct) normalized to the housekeeping gene Actin. For cell quality control, UBB and Actin expression with Ct threshold of >20 were excluded (generally 1-3% of captured cells), as this was indicative of RNA degradation.

### Homotypic and Heterotypic GJIC Assays

GJIC was measured using a novel automated parachute assay. Recipient cells are grown to confluence in a 96-cell flat bottomed plate, and the media changed to (Phenol Red-free DMEM, sodium pyruvate and 5%FBS – Assay Media) immediately before the assay. Donor cells in separate wells are incubated for 20 mins with 10uM calcein AM (Invitrogen), a membrane permeable dye that on cleavage by intracellular esterases becomes membrane impermeable, but permeable to gap junctions. After washing, trypsinization and addition of assay media, ~2500 calcein-labeled donor cells per well are dropped (‘parachuted’) onto the recipient cell layer, and calcein transfer between donor and recipient cells observed by fluorescent microscopic imaging (**Fig. 3A**). For homotypic interactions, ESCs, EECs or LP9 donor cells were parachuted onto recipient cells of the same type. For heterotypic GJIC assays, ESCs or EECs were parachuted onto LP9 recipient cells. Fluorescent, bright field and digital phase contrast images of 10-15 fields per well were captured on an Operetta automated microscope (Perkin Elmer) at 30 min intervals for approximately 2 hours. A program (developed in consultation with Perkin Elmer) allowed identification of all cells on the plate, (from phase contrast image), original donors (5-15 per field), and dye-filled recipients (based on calcein intensity – see **Fig. 3B**). Data are expressed as # of fluorescent recipient cells/# of donor cells for each condition (R/D ratio), plotted over time, and a linear regression line drawn through the data, with the slope used as a measure of coupling (**Fig. 3C**) and regression coefficient (mostly >0.8) used as a measure of assay reliability.

### Trans-mesothelial Invasion Assay

The 3-D invasion assay modeling trans-mesothelial invasion (**Fig. 5A**) has been described previously (De La Garza et.al., 2012, Ferreira *et al*., 2008, Nair *et al*., 2008). Briefly, LP9 peritoneal mesothelial cells (PMCs) were grown to confluence in 24-well invasion chamber inserts containing growth-factor-reduced Matrigel^™^, coated on 8-μm pore membranes (Corning, NY). ESCs or EECs were then labeled with CellTracker Green^®^ or DiO (Invitrogen), trypsinized and counted, prior to dropping onto the confluent layer of LP9 PMCs in the prepared inserts (~20,000 cells per insert), at which time the media below the insert was changed to the appropriate media for the invading cell (ESC or EEC). After 24 hr. incubation, non-invading cells on the upper surface of the insert were mechanically removed. Invading cells on the bottom of the membrane insert, were stained with DAPI, and 10 fields counted using an Inverted Nikon 2000 fluorescence microscope with 20x objective. Invasion assays for each cell type were performed in triplicates.

To test the role of gap junctions in the invasive process, we initially pretreated both the monolayer and dropped cells for 24 hours with 300uM GAP27 (Zealand Pharma, Copenhagen, Denmark), a peptidomimetic of the extracellular loops of Cx43 which competitively blocks Cx43 gap junction formation. Due to difficulties in obtaining consistently active peptide preparations, we subsequently shifted to 24 hour pre-treatment of the LP9 monolayer with a combination of two siRNAs to Cx43 (10 pmoles/well or 5nM final concentration) - Ambion^™^ Silencer^™^ Select) in Opti MEM (Gibco, NY) with RNAi MAX (1/100 dilution, Invitrogen), diluted 1:1 with assay media, per manufacturer’s instructions. As the siRNA transfections done immediately prior to invasion could impact cell behavior, we also knocked-down Cx43 expression using 3 different shRNAs (Dharmacon, UK) introduced into either Endometrial or LP9 Mesothelial cells via Lentivirus (carrying an RFP reported driven by a separate promoter) infection. Average infection efficiency was 58% (range 30-80%). In assessing invasion, cells expressing shRNA were identifiable as RFP positive, while unlabeled cells in the same experiment served as internal controls.

### AFM measurements of cell-cell adhesion and mesothelial integrity

We applied a Nanoscope Catalyst AFM (Bruker) interfaced with an epifluorescent inverted microscope Eclipse Ti (Nikon). AFM images were acquired with the Peak Force Quantitative Nanomechanical Mapping (QNM) mode with cells immersed in appropriate culture media. ScanAsyst probes (Bruker) with the nominal spring constant 0.4 N/m were used for imaging. The exact spring constant for each probe was determined with the thermal noise method (Butt and Jaschke, 1995). For each cell culture dish, at least 5 fields 100 by 100 μm were collected with the Peak Force set point of 2nN, and electronic resolution of 256 by 256 pixels. Nanomechanical data were processed with Nanoscope Analysis software v.1.7 (Bruker) using retrace images. Then, the Sneddon model (Sneddon, 1965) and the rules established by Sokolov (Sokolov and Dokukin, 2014) were applied for calculations of mechanical parameters.

#### Cell to cell adhesion

We attached a tester cell to a tipless probe MLCT-O10 (Bruker, cantilever A, spring constant 0.07N/m) using polyethyleneimine (PEI) as a glue (Friedrichs et. al., 2013) (**Fig. 4**). Briefly, the probes were immersed in 0.01% PEI in water for 30 min. Tester cells attachment to a culture dish was weaken by replacement of the culture medium with a non-enzymatic cell dissociation solution (Millipore) for 15-30 min in a cell culture incubator (37°C, 5%CO). Next, a single tester cell loosely attached to a culture dish was attached to a PEI covered cantilever by pressing it at 1 nN for 5-10 min. After visual inspection of successful cell attachment, the tester cell was lifted and transferred to a dish containing single tested cells. Then the tester cell was positioned over a tested cell and the cantilever slowly lowered till detection of cell-cell interactions with a force plot. The cells were left interacting for 30 to 180 sec at forces 0.5 to 5 nN and then the tester cell was lifted. During this step a force plot was recorded and the collected data applied to calculate cell – cell adhesion parameters. The force plots were baseline corrected and a maximum of adhesion between cells during their detachment was calculated (units of force, Newton) (Taubenberger, Hutmacher, and Muller 2014; Dufrêne et al. 2017).

#### Integrity of LP9 layer

LP9 cells were grown to confluence in a 60mm culture dish (**Fig. 7**). ESC cells grown in separate wells were stained with DiO, suspended, and dropped on to the LP9 monolayer at either a 1:50 or 1:20 ratio to the LP9 cells. Three hours later the cells were imaged by AFM. To calculate a tip penetration depth, cell boundaries were identified using images collected by the peak force error (PFE) channel. To exclude gap areas between cells or areas of cells growing in multilayers, PFE images were overlaid with height channel images after processing them with the flatten function of 1^st^ order. Tip penetration was calculated based on a height histogram of all data points using a difference between the prevalent maximum of cell monolayer height and the prevalent maximum depth between cells accessible for the tip.

### Data and Statistical Analysis

For microfluidic PCR analysis, cycle thresholds (Ct) were used to calculate gene expression relative to the housekeeping (HK) gene *ACTIN*. Relative expression was calculated by the formula 2^-ΔCt^, where ΔCt = Ct_gene_-Ct_HK_. Heat maps were generated for data visualization by MeV (http://mev.tm4.org/). In addition, violin plots were used to visualize single-cell expression distribution. To statistically assess this distribution between samples, Duncan’s multiple range test by R package was used to compare expression in different patient samples. For statistical analysis of cell assays, the Shapiro-Wilk normality test was first performed. t-test and ANOVA (with Holm-Sidak post-hoc analysis) were used as parametric assays after normality of data was ensured. In cases were normality test failed, Mann-Whitney rank sum test was used as a nonparametric analysis method. P values ≤ 0.05 were considered statistically significant. As data on GJIC and invasion of ESC and EEC populations from patients was normally distributed, comparisons utilized two tailed student t-tests, with a cut-off off of p < 0.05 (degrees of freedom ranged from 8 – 20). Statistical tests of all AFM data were performed and corresponding graphs prepared with OriginPro 2000b (Origin Lab). All data showed normal distribution by Shapiro-Wilk, except the 170S-LP9 (Cx43) data set in Fig. 7F, which did pass the Kolgomorov test of normality. P values were corrected using the more stringent Shapiro method.

### Cell Cluster Analysis

Single cells were clustered based on the normalized expression values using an in-house graph-based community discovery algorithm. Briefly, the algorithm starts by computing the Euclidean distances between pairs of cells, and then constructs k nearest neighbor graph, where each cell is connected to k cells that are closest to it. The best k is chosen by the algorithm with a topologically inspired criterion (Ruan, 2009). Finally, a community discovery algorithm optimizing the well-known modularity function is applied to find dense subgraphs as cell clusters (Ruan and Zhang, 2008). Note that the number of clusters is determined automatically during the optimization process.

## ACKNOWLEDGEMENTS

We would like to express gratitude to the members of the members of the High-Thoughput Screening Facility who helped with the assessment of GJIC, particularly Srikanth Polusani, the members of the clinical teams who helped in collection of patient samples, particularly Jessica Perry at UTHealth SA, and Peter Binkley curator of the UTHealth SA Ob/Gyn tissue bank. We would also like to thank Taryn Olivas and Taylor Williams who made critical contributions to the early characterization of patient samples. We would like to also acknowledge the CPRIT-funded Bioanalytics and Single-Cell Core (BASiC) facility at the University of Texas Health San Antonio for C1 cell capture and microfluidic PCR analysis and the High-throughput Screening Facility (HTSF) in the CPRIT-funded Center for Innovative Drug Discovery (CIDD). Thanks to Mr. Nicholas Lucio at the BASiC facility for his assistance with running the C1 system, and Dr. Srikanth Polusani for running the Perkin Elmer Operetta system in the HTSF.

## AUTHOR CONTRIBUTIONS

CWC and JC performed all functional experiments on cell coupling and invasion, isolated cells, and prepared them for single cell analysis; CWW, CMW, YTH and LLL conducted single cell analyses, and LLL and YTH performed statistical analyses; MJH designed and supervised the automated system for analysis of GJIC; JR performed all cell clustering analysis; LG isolated patient cells; ROB, RSS and RDR collected patient samples for analysis; MG and PO conducted all AFM experiments and analyses; NBK and BJN conceived and designed all experiments, coordinated patient sample collections and wrote the manuscript.

## SUPPLEMENTARY MATERIAL

**TABLE S1:** Genes in gene panel microarray

| Group | Gene | Protein | Group | Gene | Protein |
| --- | --- | --- | --- | --- | --- |
| **GJs** | GJA1 | Cx43 | **Kinases** | MAPK1 | ERK1 |
|  | GJA3 | Cx46 |  | MAPK3 | ERK3 |
| | GJA5 | Cx40 | | PRKACA | Prot. Kin.Aα (cat. su) |
| | GJA8 | Cx50 | | PRKACB | Prot. Kin.Aβ (cat. su) |
| | GJA9 | Cx59 | | PRKACG | Prot. Kin. Aγ (cat. su) |
| | GJB1 | Cx32 | | PRKCA | Prot. Kinase Cα |
| | GJB2 | Cx26 | | PRKCB | Prot.Kinase Cβ |
|  | GJB3 | Cx31 |  | CDK1 | Cyclin Dep.kinase1 |
| | GJB4 | Cx30.3 | | CSNK 1D | Casein Kinase 1Δ |
|  | GJB5 | Cx31.1 |  | SRC | c-src |
|  | GJB6 | Cx30 |  | NOV | CCN3 |
|  | GJB7 | Cx25 |  | PLCB1 | Phosholipase B1 |
|  | GJC2 | Cx47 |  |  |  |
| **Adhesion/TJs.** | TJAP1 | TJ assoc. protein | **Markers** | VIM | Vimentin |
|  | TJP1 | ZO1 |  | MME/CD10 | Metallo-endopept. |
|  | TJP2 | ZO2 |  | EPCAM | Ep CAM |
|  | CAV1 | caveolin |  | KRT 18 | Cytokeratin 18 |
|  | CDH2 | N-cadherin | **Stand.** | GAPDH | GAP dehydrog. |
|  | SNAIL1 | Snail 1 |  | ACTIN-B | Actin |
|  | CTNNB1 | beta-catenin |  |  |  |
|  | DBN1 | Drebin 1 |  |  |  |

**Figure S1:**
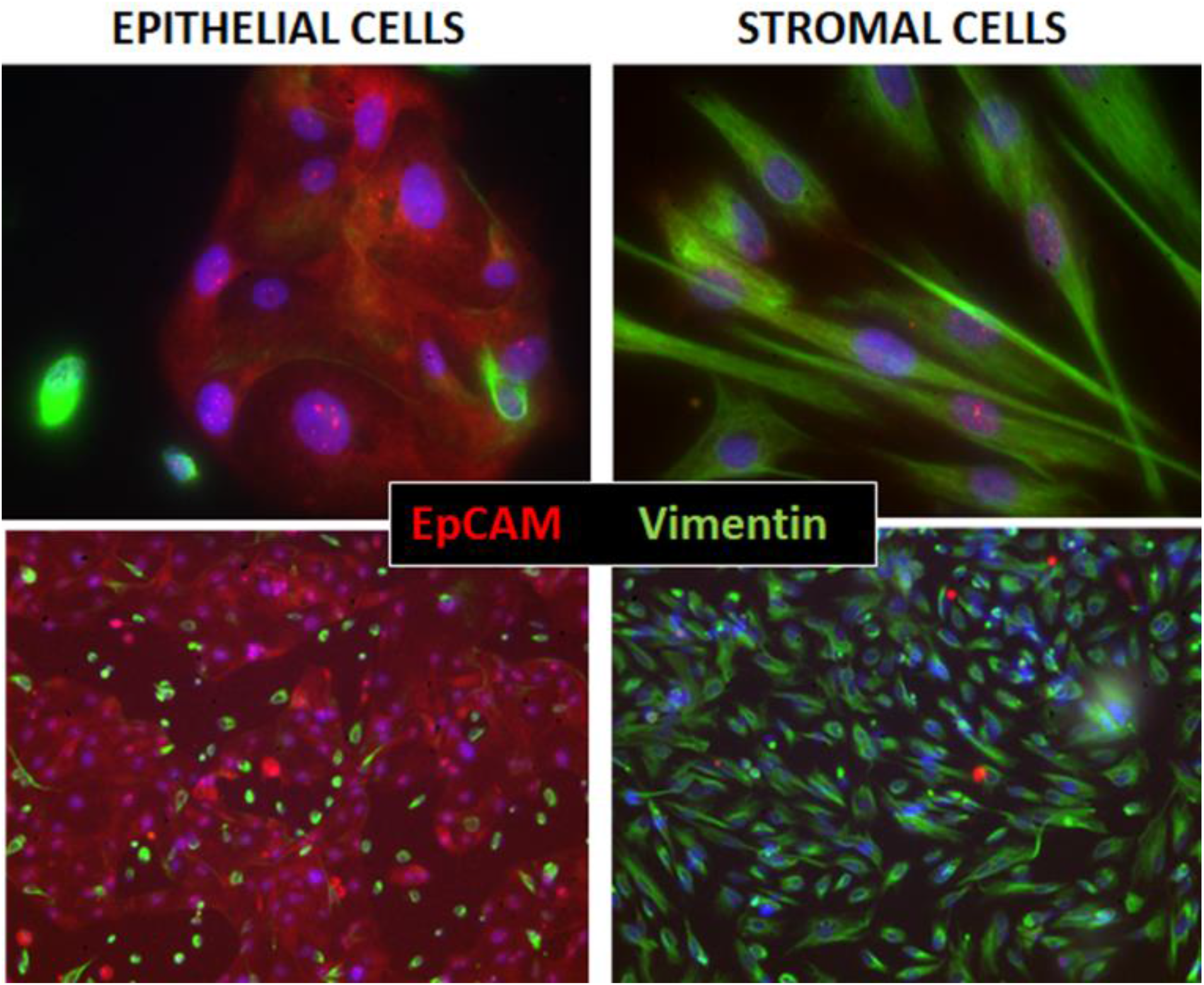
Immunocytochemical assessment of Epithelial and Stromal cell isolations from two patients. Endometrial cells are separated by differential sedimentation and adhesion into epithelial (left) and stromal populations (right). Immunohistochemistry [40x (top) and 10x (bottom)] demonstrate the purity of the isolations by dual staining with EpCAM antibodies to mark epithelial cells (red), and Vimentin antibodies to mark stromal cells (green). Nuclei are stained blue with DAPI in both.

**Figure S2:**
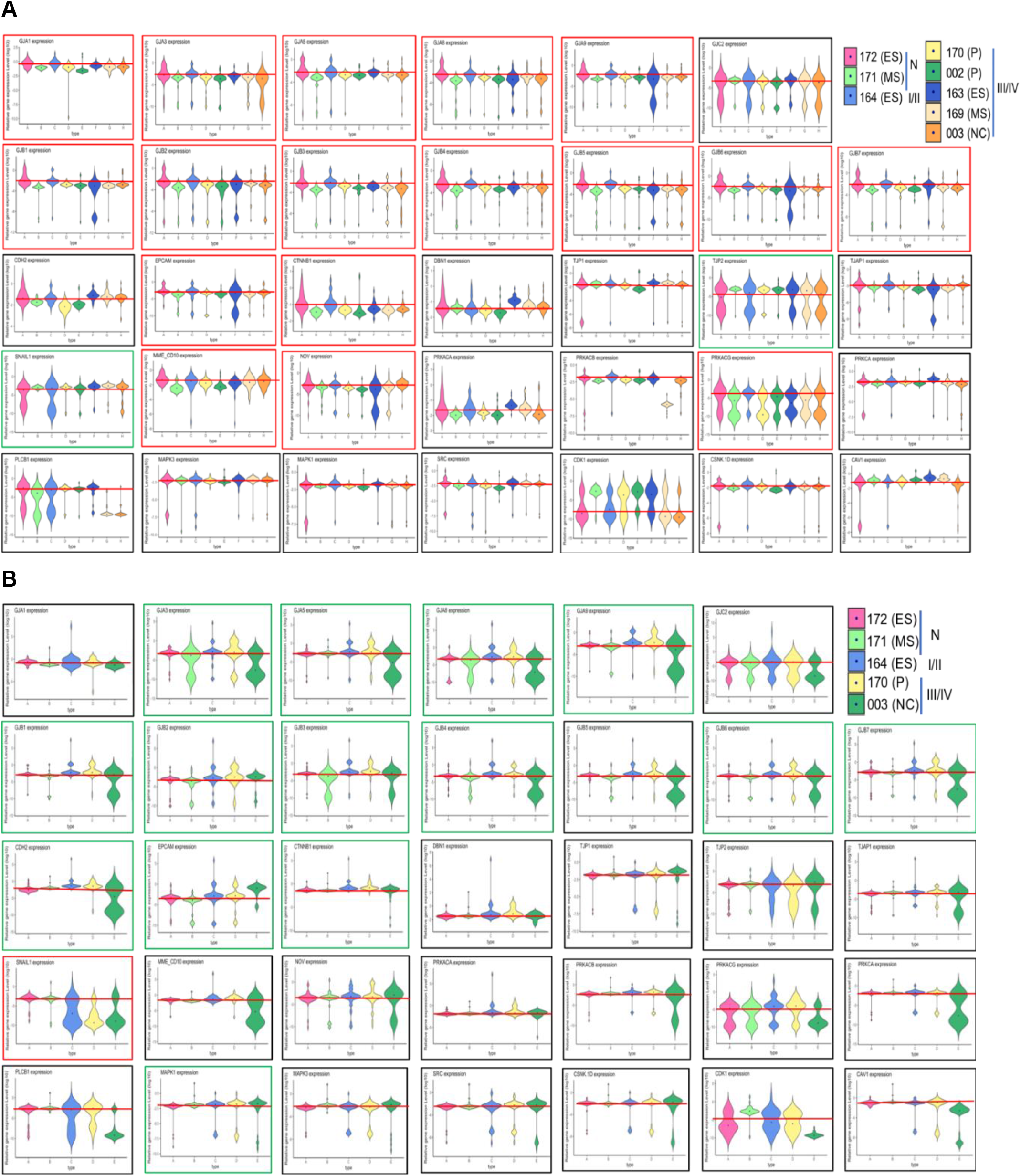
Violin plots of average cell expression levels for all genes for which signals were detected and for all patients in stromal(A) and Epithelial cells (B). Patients are color coded in legend, and are arranged in each plot from left to right as control (N), early stage (I/II) and late stage endometriosis. Phase of the cell cycle when samples were collected are indicated in the legend (P = proliferative; ES = early secretory; MS = mid-secretory; NC = non-cycling). The average value of control and early stage endometriosis in ESCs (**A**) or just control samples in EECs (**B**) is shown as a red horizontal line for reference, and the general pattern with endometriosis is indicated by arrows, and color of margin (red = decreased expression with endometriosis, green = increased expression). Note that in ESC samples collected at the MS stage and EECs (and in a few cases ESCs) collected from non-cycling patients, some genes show a divergence from general pattern of gene expression, indicating effects of the hormonal environment on expression.

